# miR-424(322) is a molecular switch controlling pro-inflammatory vs anti-inflammatory skin DC subset differentiation by modulating TGF-β signaling

**DOI:** 10.1101/2020.09.07.285627

**Authors:** Victoria Zyulina, Koon-Kiu Yan, Bensheng Ju, Christina Passegger, Carmen Tam-Amersdorfer, Qingfei Pan, Tommaso Sconocchia, Christian Pollack, Bridget Shaner, Armin Zebisch, John Easton, Jiyang Yu, Jose M. Silva, Herbert Strobl

**Author notes:** Corresponding authors: Herbert Strobl MD, Otto Loewi Research Center, Division of Immunology and Pathophysiology, Heinrichstrasse 31A, 8010 Graz, Austria,;, Jose M.Silva PhD, Department of Pathology, Icahn School of Medicine at Mount Sinai, Gustave L. Levi PI., NY, NY 10029, USA.

## Abstract

TGF-β family ligands are key regulators of dendritic cell (DC) differentiation and activation. Epidermal Langerhans cells (LCs) require TGF- β family signaling for their differentiation and canonical TGF-β1 signaling secures a non-activated LC state. LCs reportedly control skin inflammation and are replenished from peripheral blood monocytes, which also give rise to pro-inflammatory monocyte-derived DCs (moDCs). Among all the miRNAs differentially expressed in LC vs moDCs, we observed miR-424 to be strongly induced during moDC differentiation from monocytes. We discovered that miR-424 is required for moDC differentiation from human and murine precursor cells *in vitro* and for inflammation-associated moDC *in vivo*. Mechanistically we found that low levels of miR-424 facilitate TGF-β1-dependent LC differentiation at the expense of moDC differentiation. Loss of miR-424 in monocyte/DC precursors resulted in the induction of TGF-β pathway. Therefore, miR-424 plays a decisive role in anti-inflammatory LC vs pro-inflammatory moDC differentiation from monocytes, and its repression allows TGF-β ligands to promote LC differentiation.

**Short summary:** Monocytes give rise to two distinct DC subsets in skin inflammation, exhibiting opposite roles in inflammation. This study identified miR-424(322) as a molecular switch controlling pro-inflammatory (moDC) vs anti-inflammatory LC subset differentiation by modulating TGF-β signaling.

## Introduction

The mononuclear phagocyte and dendritic cell (DC) system comprises a large number of cell subsets that markedly differ in function and anatomic location. While many of these subsets are known to populate tissues in the steady-state, monocyte-derived DCs (moDCs) are predominantly associated with inflammatory lesions such as psoriatic cutaneous manifestations (Wohn et al., 2013). Human moDCs can be generated *in vitro* from monocytes in response to GM-CSF plus IL-4 (Sallusto & Lanzavecchia, 1994) and are phenotypically similar to moDCs in inflammatory skin lesions in psoriasis and atopic dermatitis patients (Kraft et al., 2002; Rescigno & Di Sabatino, 2009; Rimoldi et al., 2005). Another skin-associated DC subset, i.e. epidermal Langerhans cells (LCs), is known to counteract and limit skin inflammation in chronic psoriatic lesions (Glitzner et al., 2014). LCs form dense cellular networks in basal/suprabasal epithelial layers that are established prenatally from embryonic and fetal precursors and can also be replenished from circulating monocytes in response to injury and inflammation (Honda, Egawa, & Kabashima, 2019). Therefore, monocytes can give rise to two DC subsets with contrasting functions in skin inflammation, i.e. anti-inflammatory LCs vs pro-inflammatory moDCs. However, little is still known about the intracellular mechanism underlying LC vs moDC differentiation from monocytes.

Monocyte committed progenitor cells (moPs) originate from shared progenitor cells with conventional DCs and plasmacytoid DCs (Merad, Sathe, Helft, Miller, & Mortha, 2013; Naik et al., 2007) (Geissmann et al., 2010). Further differentiation of moPs into monocytes/macrophages is induced by the master transcription factor Kruppel-like factor KLF4(Feinberg et al., 2007; Naik et al., 2007). Sub-lineage commitment of monocytes into moDCs occurs at the expense of macrophage differentiation and is induced by the transcription factors AhR and IRF4 while being repressed by MafB (Goudot et al., 2017). Conversely, monocyte to LC differentiation depends on the induction of the master positive transcriptional regulator of LCs, RUNX3 (Fainaru et al., 2004), which is transcriptionally induced by PU.1 (Chopin et al., 2013), and requires TGF-β1 signaling in precursors undergoing LC differentiation (Jurkin et al., 2017). RUNX3 gene expression is negatively regulated by KLF4 which in turn is repressed by Notch signaling within the skin, thus enabling LC differentiation from monocytes (Jurkin et al., 2017). Despite this current knowledge on transcription factors controlling moDC vs LC differentiation, how extracellular signal input in monocytes is relayed to induce downstream transcriptional programs leading to LC vs moDC fate decisions still remained poorly understood.

Given that LCs vs other DCs are differentially affected by deletion of Dicer, the key enzyme in miRNA processing (Kuipers, Schnorfeil, Fehling, Bartels, & Brocker, 2010), we rationalized that miRNAs might represent switch factors involved in binary LC vs moDC fate decisions of monocytes. Thus, we previously searched for miRNAs inversely expressed by LCs vs moDCs generated *in vitro* from human monocytic cells (Jurkin et al., 2010, Lim et al., 2020). Among several differentially expressed miRNAs, miR-424 stood out as the most differentially expressed miRNA when comparing moDCs vs LCs(Jurkin et al., 2010). Interestingly, miR-424 was shown to promote monocyte/macrophage differentiation in human HL-60 cells (Kasashima, Nakamura, & Kozu, 2004) and CD34^+^ hematopoietic progenitor cells (Rosa et al., 2007). However, a potential role of miR-424 in LC vs moDC subset differentiation remained unexplored and its *in vivo* role in myelopoiesis has not been addressed thus far.

Using human and murine models, we here showed that miR-424 is induced during moDC differentiation from monocytes and its upregulation is required for inflammation-linked moDC differentiation. In contrast, low miR-424 expression facilitates the differentiation and accumulation of non-activated LCs, associated with augmented expression of genes of the TGF-β family.

## Results

### MiR-424 is induced during moDC and repressed during LC differentiation of monocytic cells

Given the importance of the miRNA pathway in DC differentiation *in vivo* (Kuipers et al., 2010), we previously screened purified fractions of *in vitro* generated LCs and moDCs for differentially expressed miRNAs. This led to the identification of miR-424 to be 6 times higher expressed in moDCs than in LCs(Jurkin et al., 2010). The miR-424 is expressed as a cluster with the miR-503 from a locus located in the X chromosome (Forrest et al., 2010). miR-424 and miR-503 belong to the same microRNA family, share seed sequence and target profile (Forrest et al., 2010). Interestingly, it has been reported that miR-503 has a much lower expression level (~100 times) than miR-424 (Llobet-Navas, Rodriguez-Barrueco, Castro et al., 2014). Thus, we first evaluated the expression of miR-503 in our samples. As expected, miR-503 showed the same pattern of expression as miR-424 and its expression was 3 times higher in moDC than in LCs; however, its expression compared to miR-424 was very low (Suppl. Fig. 1A). Because of the low expression of the miR-503, together with previous findings showing that miR-424 plays a non-redundant functional role in human monocyte/macrophage differentiation (Kasashima et al., 2004; Rosa et al., 2007), we decided to focus on miR-424 in subsequent experiments.

To study the role of miR-424 in DC subset specification we employed a previously described *in vitro* differentiation model of human LCs vs moDC differentiation from monocyte intermediates generated from CD34^+^ hematopoietic progenitor cells (Jurkin et al., 2010). Briefly, a combination of cytokines induces the pre-generation of monocytic precursor cells from progenitor cells, which then can be induced to differentiate into CD1a^+^CD207^+^ LCs in response to GM-CSF, TNFα plus TGF-β1 or into CD1a^+^CD11b^+^CD209^+^ moDCs in response to GM-CSF plus IL-4 (Fig. 1A). Importantly, the omission of TGF-β1 from the LC generation cultures (“LC mix”) leads to abrogation of LC differentiation, in favor of induction of CD11b^+^ cells exhibiting similarities with interstitial/dermal DCs (Platzer, Jorgl, Taschner, Hocher, & Strobl, 2004). Moreover, IL-4 represses LC characteristics when added to TGF-β1-dependent LC generation cultures (Caux et al., 1999; Heinz et al., 2006).

**Figure 1.**
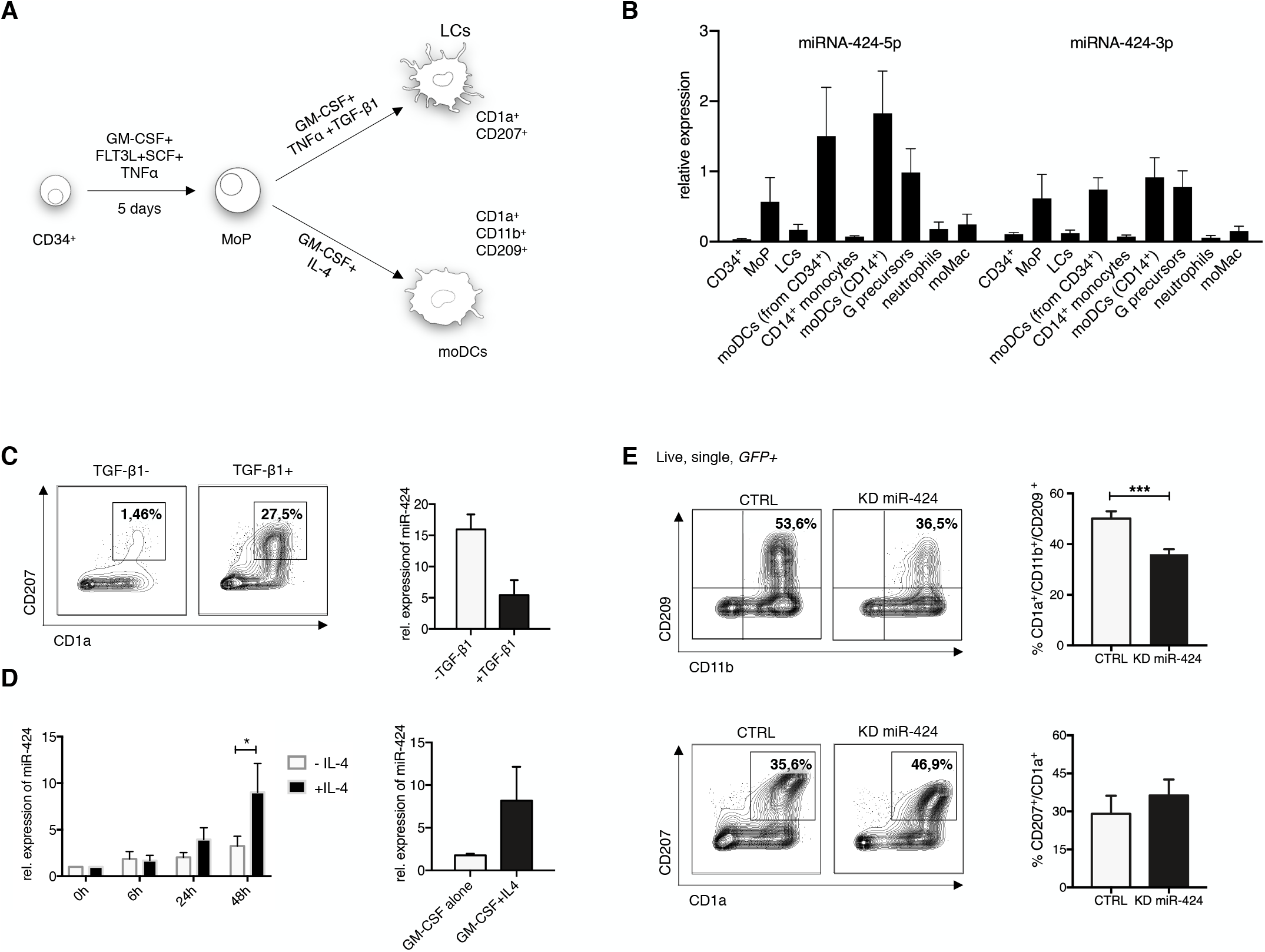
MiR-424 is required for moDCs but not LC differentiation from human CD34^+^ hematopoietic progenitor cells. **A**. Schematic representation of moDCs and LCs differentiation model from CD34^+^ cells *in vitro*. **B**. Relative expression of miR-424-5p and miR-424-3p in different myeloid cell types measured by quantitative Taqman PCR (n=3-5, ± SEM). **C**. Representative flow cytometry analysis of CD34^+^ cells differentiated for 7 days into LCs in cytokine cocktail with and without TGF-β1. Graph shows relative expression of miR-424 in + /− TGF-β1 LCs on day 7 after differentiation from CD34^+^ progenitors (n=3, ± SEM). **D**. Graph shows relative miR-424 expression in +/− IL-4 moDCs generated from MoPs (n=6, ± SEM) in the left panel and from CD14^+^ monocytes (n=3, ± SEM) in the right panel. Data were analyzed using paired 2-tailed Student’s t-test (*P<0.05). **E**. Lineage marker profile of gated GFP^+^ CD1a^+^ miR-424 knockdown, induced into moDCs or LCs differentiation under lineage specific conditions. Data represent the mean ± SEM of 3-6 donors, 2-tailed Student’s t test, *** P<0.001.

The comparison of miR-424 expression levels in different myeloid cell types showed that freshly isolated CD14^+^ blood monocytes, CD34^+^ cells and peripheral blood neutrophils expressed very low levels of miR-424, and miR-424 levels remained low during GM-CSF-induced monocyte-derived macrophage (moMac) generation (Fig. 1B). In contrast, miR-424 expression was strongly increased during moDC differentiation, irrespective of whether GM-CSF/IL-4-dependent moDCs were generated from monocytes or from CD34^+^ cells. (Fig. 1B). *In vitro* generated MoPs (schematically shown in Fig. 1A) exhibited inverse regulation of miR-424 upon their differentiation in to LCs vs moDCs (Fig. 1B), consistent with our previous data (Jurkin et al., 2010). Omission of TGF-β1 from day 7 LC generation cultures (i.e. containing GM-CSF+ TNFα, but no exogenous TGF-β1) resulted in elevated miR-424 expression levels (Fig. 1C), indicating that TGF-β1 represses miR-424 during LC lineage instruction. Next, we studied miR-424 expression regulation downstream of IL-4 during moDC differentiation (Sander et al., 2017). miR-424 expression levels gradually increased in response to IL-4 co-stimulation within 48h during moDC differentiation from MoPs (Fig.1D, left panel), similarly IL-4 promoted miR-424 upregulation during moDC differentiation from peripheral blood monocytes (Fig. 1D, right panel, and suppl. Fig. 1B). Thus, miR-424 is induced during moDC differentiation in response to IL-4.

### miR-424 is required for human moDC but not for LC differentiation

Given the observed upregulation of miR-424 during moDC differentiation and its low levels in LCs, we next studied whether miR-424 is required for moDC vs LC differentiation from CD34^+^ hematopoietic progenitor cells. Hence, lentiviral vector-mediated silencing of miR-424 was performed in CD34^+^ progenitors during the first 5 days (leading to MoP generation), and cells were then induced to differentiate into moDCs vs LCs as shown in Fig. 1A. The modulation of miR-424 expression using lentiviral vector was confirmed by qPCR (Suppl. Fig. 1C). Lentiviral vector transduced GFP^+^ cells were analyzed for marker combinations specific for moDCs (CD1a^+^CD11b^+^CD209^+^) or LCs (CD1a^+^CD207^+^). While miR-424 knockdown resulted in diminished frequencies of GM-CSF^+^IL-4-induced moDCs relative to parallel cultures of control transduced cells (Fig. 1E, upper panel), GM-CSF^+^TNFα ^+^TGF-β1-dependent LC differentiation remained unimpaired; instead on average slightly higher percentages of LCs were observed in response to miR-424 knockdown (Fig. 1E, lower panel). Taken together, our data shows that miR-424 is induced during moDC differentiation and promotes moDC differentiation. Inversely, endogenous miR-424 is repressed during LC differentiation and miR-424 knockdown failed to impair LC differentiation.

### miR-424(322) is required for moDC differentiation *in vivo*

To investigate the role of miR-424 in moDC differentiation in a tractable *in vivo* model we turned our studies to a miR-knock-out mouse model. For this we used the miR-424(322)/503 model. miR-322 is the murine orthologue of the human miR-424. Here, it will be called miR-KO for simplification (Llobet-Navas, Rodriguez-Barrueco, de la Iglesia-Vicente et al., 2014). First, we induced psoriasis-like inflammation in miR-KO vs WT mice by topical application of TLR7 agonist Imiquimod (IMQ) during 6 consecutive days (Che et al., 2019; Rodriguez-Barrueco et al., 2017; Wang et al., 2019). Then, we analyzed the frequencies of Ly6C^int^MHCII^high^ (T3) and Ly6C^low^MHCII^high^(T4) moDCs in the skin of IMQ-treated miR-KO mice vs WT mice (Fig. 2A). Expectedly, WT mice showed a substantial increase in frequencies of moDCs (Fig. 2B). Consistent with our human *in vitro* findings, we observed that moDC subsets were diminished in the dermis of miR-KO relative to WT mice upon inflammation. In addition to exhibiting lower frequencies of moDCs, miR-KO mice also exhibited reduced frequencies of macrophages (T1) in the inflamed dermis (Suppl. Fig. 2A). Conversely, there was no detectable difference in the numbers of moDCs in the steady-state (Fig. 2B). Moreover, frequencies of CD103^+^ cDCs, CD11b^+^ cDCs and Ly6C^high^ monocytes(T2) remained equivalent in KO-mice vs WT controls (Suppl. Fig. 2B). Overall, these results indicate that miR-424(322) promotes murine moDC (T3+T4) differentiation. MoDCs are known as the principal contributors to skin inflammation and DC depletion prevents IMQ-induced skin inflammation(Singh et al., 2016). Consistently, IMQ treated miR-KO mice displayed significantly attenuated ear swelling relative to wild-type (WT) mice (Fig. 2C left panel), confirmed by microscopical measurements of the epidermis at day 7 (Fig. 2C, middle and right panel). In line with changes in local inflammation, there was no difference in body weight loss between the groups (Suppl. Fig. 2C). In conclusion, miR-KO mice displayed decreased frequencies of moDCs associated with diminished psoriatic-like skin inflammation relative to WT mice.

**Figure 2.**
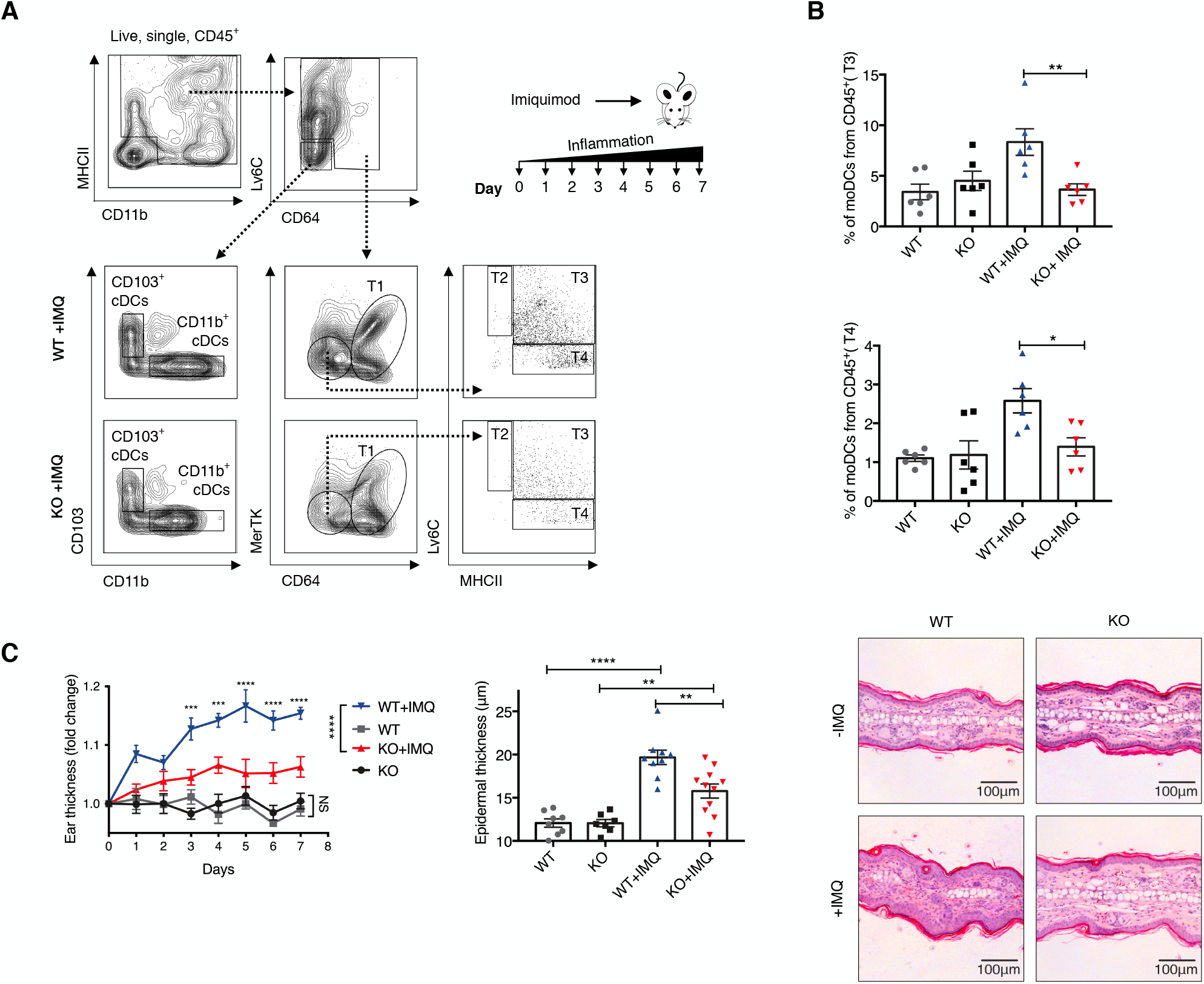
Mir-424(322) is required for moDCs differentiation *in vivo*. **A**. Gating strategy for cell subset identification in the dermis of mouse ears on day 7 of experiment: macrophages (T1), monocytes (T2), Ly6C^++^ moDCs (T3), Ly6C^+^ moDCs (T4). Schematic overview of IMQ-induced psoriasis-like skin inflammation in mice. **B**. Graph represents percentage of phenotypically defined moDCs subsets T3 and T4 in mouse dermis quantified by flow cytometry on day 7 of experiment (n=6, ± SEM, unpaired Student’s t test, *P<0.05, **P<0.01). **C**. Ear swelling of both mouse ears in WT and miR-424 KO mice measured on the indicated days (n=6-8 mice per group, ± SEM), two-way Anova, *** P<0.001, **** P<0.0001. Epidermal thickness of each mouse measured on H&E staining on experimental day 7 (n=4 mice per group, ± SEM, unpaired Student’s t test, **P<0.01, **** P<0.0001). Each dot represents a mean of 4-5 measurements. The right panel shows representative H&E staining picture of the mouse ear.

### miR-424(322) deficient mice exhibit elevated numbers of non-activated LCs

As demonstrated above (Fig. 1B), miR-424 expression increased during moDC differentiation but is repressed during LC differentiation from MoPs. Our human differentiation culture model system mimics moDC vs LC development from monocytes during inflammation. Thus, we next investigated how the absence of miR-424(322) influences LC frequencies in inflammatory lesions *in vivo*. Immunohistology of epidermal sheets prepared from untreated mice revealed an undisturbed network of LCs, with on average slightly increased frequencies of CD207^+^ LCs in miR-KO mice relative to WT control (Fig. 3A). FACS analysis of mouse epidermal sheets revealed a minor portion of LCs from WT mice expressing CD86 in the steady-state, and percentages of CD86^+^ cells expectedly increased after IMQ treatment (Fig. 3B and C). Interestingly, LCs from miR-KO mice exhibited reduced percentages CD86^+^ cells compared to WT mice in steady-state and after IMQ treatment (Fig. 3B and C), indicative of a less activated LC phenotype. Upon activation, LCs emigrate from the epidermis to the skin-draining lymph nodes. Frequencies of LCs among CD45^+^ cells in draining lymph nodes showed tendency to decrease in IMQ treated miR-KO mice compared to WT mice (p value=0.0952, Suppl. Fig. 2D).

**Figure 3.**
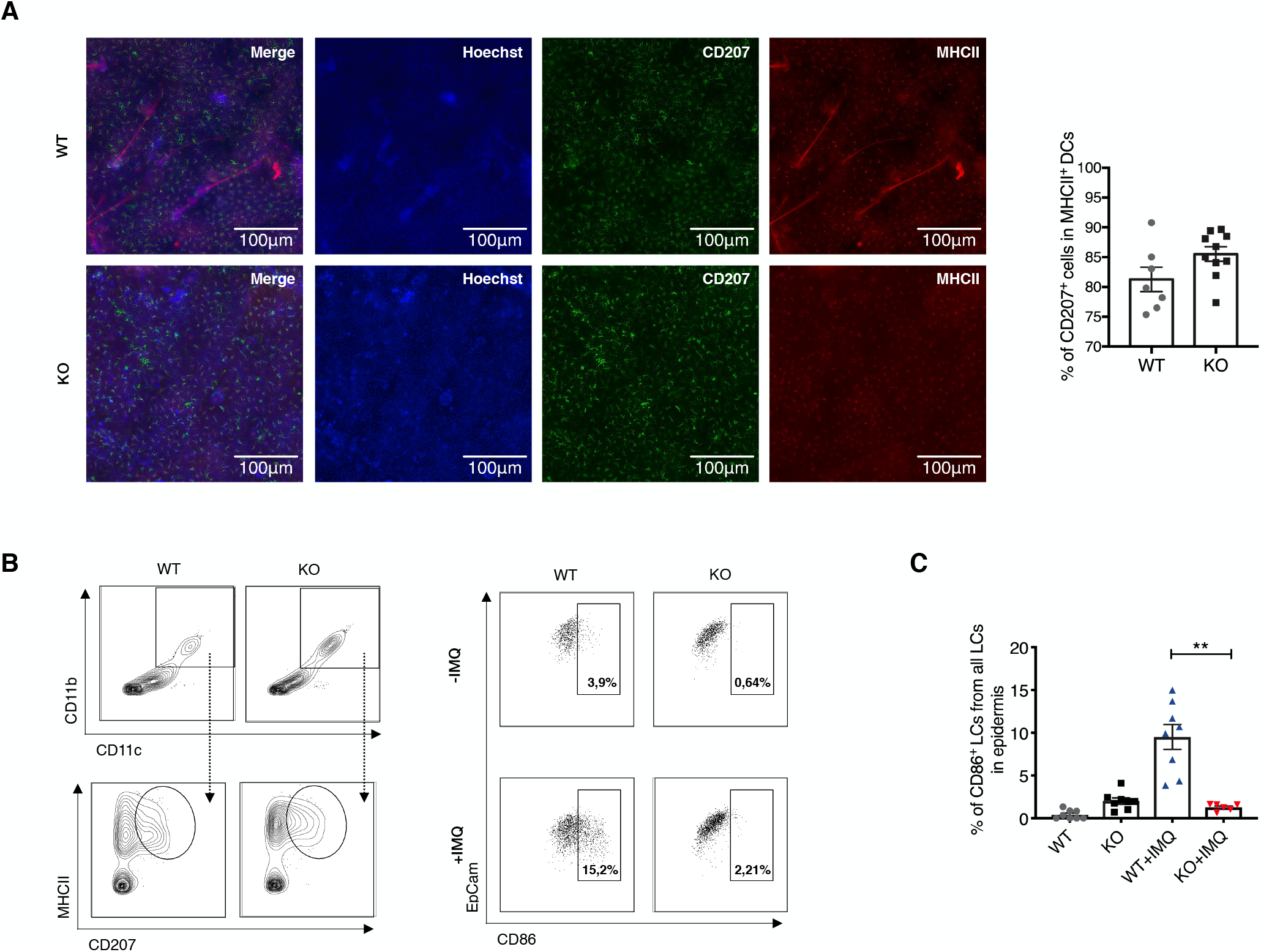
miR-424(322) deficient mice exhibit on average slightly elevated numbers of non-activated LCs. **A**. Mouse epidermal sheets staining for CD207^+^ and MHCII^+^ DCs in steady-state. Frequency of CD207 LCs in MHCII DCs in immunofluorescent staining of mouse epidermal sheets on day 7 of IMQ experiment. Each symbol represents one mouse epidermal sheet. **B**. Gating strategy for mouse LCs in the epidermis of the ear on day 7 of IMQ experiment. **C**. Graph shows frequency of CD86^+^ LCs in the epidermis of mouse ear (n=8 mice per group, ± SEM, unpaired Student’s t test, **P<0.01).

### moDC precursors in the bone marrow are present at equivalent frequencies in WT and miR-424(322)-deficient mice

Inflammation-associated moDCs develop from bone marrow (BM) precursor cells. Thus, it was possible that the diminished frequencies of moDCs observed *in vivo* in IMQ treated miR-424(322)-deficient mice are mediated by defects in their BM precursors. To evaluate this possibility, we phenotypically analysed freshly isolated BM cells of WT vs miR-KO mice to identify precursors of moDCs (lin^−^CD115^+^Ly6C^+^FLT3^−^CD11c^−^; i.e. R1 cells, Fig. 4A) (Menezes et al., 2016). Murine Ly6C^+^ monocytes comprise a heterogeneous population and give rise to iNOS^+^ macrophages (R1), inflammatory moDCs upon GM-CSF exposure (R2) or FLT3 dependent DCs (R3)(Menezes et al., 2016). To investigate whether the numbers of distinct monocytic precursors are impaired in miR-KO mice, we quantified all three populations in freshly isolated mouse BM. (Fig. 4A). BM from WT and miR-KO mice exhibited equivalent frequencies of R1, R2 and R3 cells (Fig. 4B). Therefore, previously described BM precursors of moDC are present at equivalent frequencies in miR-KO vs WT mice.

**Figure 4.**
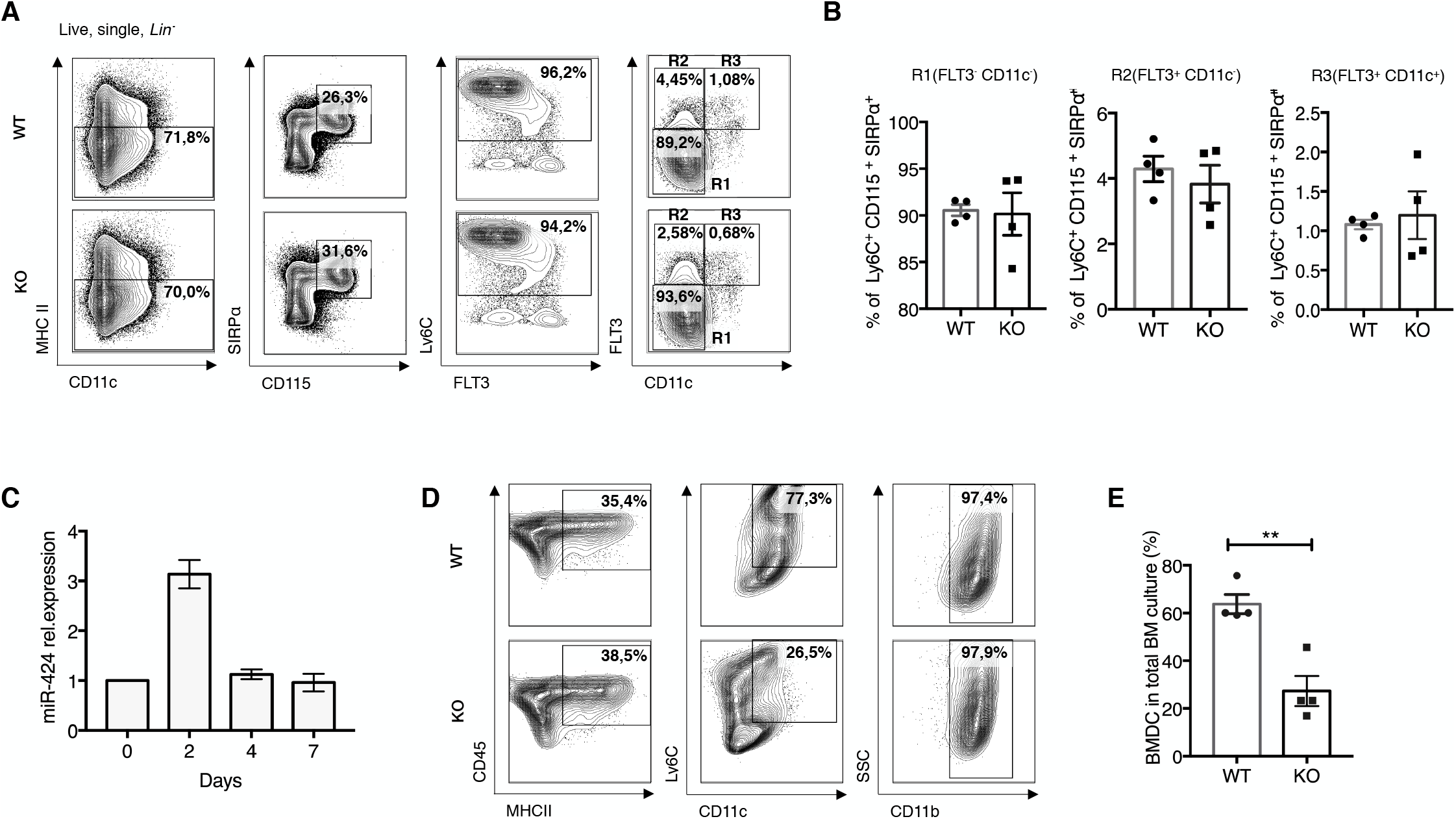
miR-424(322) is essential for *ex vivo* BMDC differentiation but it does not interfere with monocytic precursors in mouse bone marrow. **A**. Flow cytometry analysis of fresh isolated mouse bone marrow for dendritic cell precursors. **B**. Graph depicts frequency of R1 (precursors for iNOS macrophages), R2 (moDCs precursors), R3 (pre-DC precursors) from Ly6C^+^CD115^+^SIRPα^+^ population. **C**. MiR-424(322) endogenous levels in BMDCs cultures over time. **D**. BM cells were cultured with GM-CSF for 7 days. Flow cytometry analysis of CD11c, Ly6C and CD11b expression on CD45^+^MHCII^+^ cells. **E**. Percentage of BMDCs in GM-CSF BM cultures. Data represent ± SEM of 4 mice, 2-tailed Student’s t test, ** P<0.01;

### Altered gene expression profile in GM-CSF stimulated miR424(322)−/− vs WT bone-marrow cells

To identify genes and pathways differentially expressed and activated between WT vs miR-KO immune cells we compared their expression profiles during differentiation into DC subsets. GM-CSF-stimulated bulk cultures of murine BM cells give rise to several (sub)lineages of monocyte/macrophage and DC subsets (Helft et al., 2015). Monitoring of miR-424(322) expression levels revealed a transient upregulation of miR-424(322) peaking at 48h after GM-CSF stimulation (Fig. 4C). Given the above-described observations of undisturbed frequencies of previously characterized monocyte/macrophage/DC BM precursor cells in miR-KO mice, we rationalized that GM-CSF-induced BM cultures are suited to identify genes up-or downregulated in response to miR-424(322) deficiency. First, we studied whether miR-424(322) is required for GM-CSF-dependent DC differentiation. Indeed, phenotypic analysis of day 7 generated cells from miR-KO vs wild-type mice revealed diminished generation of MHCII^+^Ly6C^+^CD11c^+^CD11b^+^ BMDCs in miR-KO mice (Fig. 4D and E). We next performed RNA sequencing to identify genes differentially expressed by WT vs miR-KO BM cells early in response to GM-CSF stimulation. As expected from above-described phenotypic analyses, unstimulated BM cells from WT vs miR-KO mice on day 0 exhibited only minor differences in gene expression (Fig. 5A). After 48 hours of GM-CSF stimulation 418 genes were significantly upregulated or downregulated in WT condition (Fig. 5B). Then we determined those genes among GM-CSF response genes that are differentially expressed by WT vs miR-KO cells at 48h (Fig. 5C). Interestingly, most of these genes were substantially upregulated in miR-424(322) deficient cultures, including transcription factors (*Atf5, Mitf* and *Zfpm1*), surface receptors (Tnfrsf9, CCR4, Fas, Tlr4, Scn4a and Slc40a1) and ligands involved in cell signaling (Spp1, Mmp13, Hbegf, Prok2, Tnf, Lama5, Vcan, Bst1). 19 genes were downregulated, with some being stronger downregulated in WT (e.g. Tinagl-1, Sh3fc2); others were stronger downregulated miR-KO cells (e.g. Dntt). These data suggested that miR-424(322) induction after 48h in GM-CSF treated BM cells correlates with altered expression of GM-CSF response genes resulting in the impairment of GM-CSF-dependent moDC differentiation.

**Figure 5.**
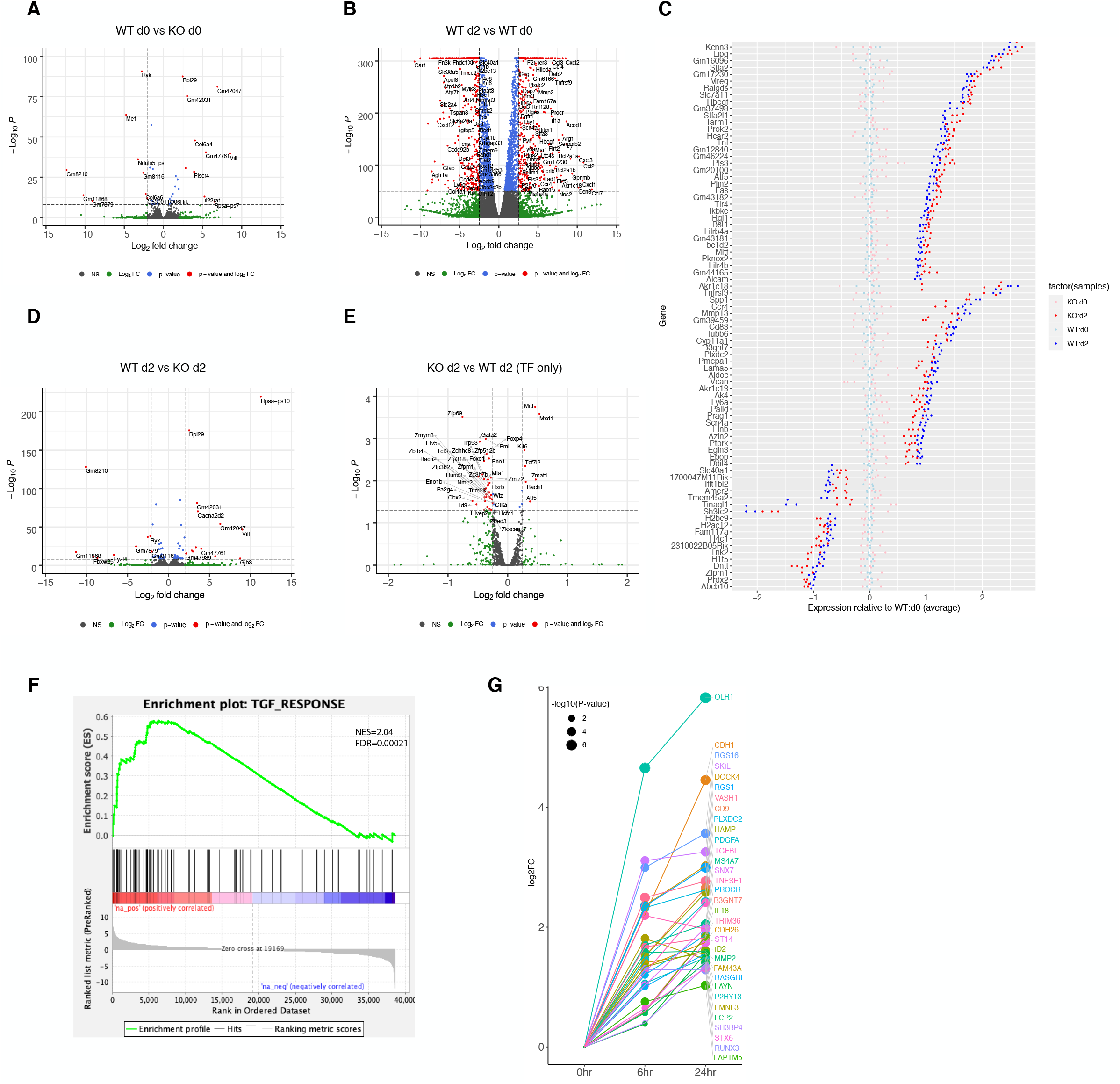
Altered gene expression profile in GM-CSF stimulated miR424(322)−/− vs WT bone-marrow cells. **A**. Volcano plot representation of differential expression analysis of genes in unstimulated BM cells from WT vs miR-KO mice (day 0). **B**. Volcano plot representation of differential expression analysis of genes in BM cells from WT mice on day 0 vs day 2 of GM-CSF stimulation. **C**. The graph shows differential gene expression between WT vs miR-KO cells at day 0 and day 2. **D**. Volcano plot representation of differential expression analysis of genes in BM cells from WT vs miR-KO mice on day 2. **E**. Volcano plot representation of differential expression of transcription factors in BM cells from WT vs miR-KO mice on day 2. **F**. Gene set enrichment analysis (GSEA) performed on comparisons of TGF-β and miR-KO signature genes. **G**. Gene expression data of TGF-β signature genes on different time points of LC differentiation overlapped with genes upregulated in miR-KO mice.

### Concomitant with LC differentiation TGF-β1-signature genes are induced in GM-CSF stimulated miR-KO BM cells

TGF-β1 signaling is known to induce LC differentiation from monocytopoietic cells at the expense of macrophage and moDC differentiation *in vitro* (Geissmann et al., 1998; Strobl et al., 1996). Since we observed impairment in moDC and macrophage differentiation in IMQ treated miR-KO mice and this effect was associated with on average elevated frequencies of non-activated epidermal LCs (Fig. 2 and 3), we expanded our studies to investigate whether TGF-β1 response genes might be induced in GM-CSF stimulated miR-KO BM cells. To address this question, we first identified 1395 genes differentially expressed on day 2 of differentiation in WT vs miR-KO BM cells (Fig. 5D), among them 47 differentially expressed transcription factors (Fig. 5E). Previously we identified TGF-β response genes in human CD34^+^CD45RA^hi^CD19^−^ progenitors committed to LC differentiation at 6 and 24 h after TGF-β1 addition (Jurkin et al., 2017). Based on these two datasets, we next performed set enrichment analysis (GSEA) to see whether loss of miR-424(322) in 48h stimulated GM-CSF treated BM cells results in the enrichment of such TGF-β1-response genes during LC commitment. This was indeed the case (Figure 5H). As can be seen from Fig. 5G, 33 genes among the previously identified TGF-β1-LC response genes were identified to be significantly induced in miR-KO vs WT cells. This list included transcriptional regulators that were previously shown to be essential for LC commitment, i.e. RUNX3 (Fainaru et al., 2004), ID2 (Hacker et al., 2003), TGF-β1 (Kaplan et al., 2007) and surface molecule CDH1/E-cadherin (Riedl et al., 2000) functionally implicated in LC differentiation (Table S1). Therefore, loss of miR-424(322) in GM-CSF stimulated BM cells leads to the induction of genes that were functionally shown to be essential for LC commitment and differentiation.

### Loss of miR-424 facilitates TGF-β1-dependent LC differentiation at the expense of moDC differentiation from human progenitor cells

To validate above findings (Fig. 5G), we next studied whether loss of miR-424 in monocytic cells indeed results in augmented TGF-β1-dependent LC differentiation. As described above, moDC differentiation of monocytic cells in response GM-CSF plus IL-4 stimulation is accompanied by the induction of miR-424 within 48h (Fig. 4C). Therefore, we studied whether lentiviral miR-424 knockdown might facilitate TGF-β1-dpendent LC differentiation from 48h GM-CSF/IL-4 pre-stimulated monocytic cells. We first evaluated consequences of IL-4 addition to LC generation cultures (“LC mix”, Fig. 6A). It was previously shown that IL-4 addition to GM-CSF/TNFα containing cultures of CD34^+^ hematopoietic progenitor cells represses TGF-β1-induced LC differentiation in favor of DCs and macrophages (Caux et al., 1999). Consistently, the addition of IL-4 represses LC differentiation in favor of CD209^+^(DC-SIGN^+^) moDC differentiation (Fig. 6A). Next we pre-treated *in vitro* generated MoPs (according to Fig. 1A) with GM-CSF plus IL4 for 48h to induce miR-424 (Fig. 1D) and moDC differentiation, followed by the addition of TGF-β1-containing LC cytokine mix. Parallel cultures were maintained with GM-CSF plus IL-4 without LC cytokine mix. CD34^+^ cells were transduced with lentiviral vectors to induce miR-424 knockdown or CTRL vector. miR-424 knockdown led to the generation of higher percentages of CD207^+^CD1a^+^ LCs and lower percentages of CD209^+^(DC-SIGN^+^) moDCs relative to control transduced cultures. Therefore, miR-424 is induced downstream of IL-4 during moDC differentiation and miR-424 functionally interferes with TGF-β1-dependent LC differentiation during moDC differentiation. This is consistent with above observation of induced TGF-β1 LC response genes in miR-KO vs WT BM cells (Fig. 5G). Moreover, these data support that miR-424 is involved in lineage fate decisions of monocytes towards LC vs moDC differentiation.

**Figure 6.**
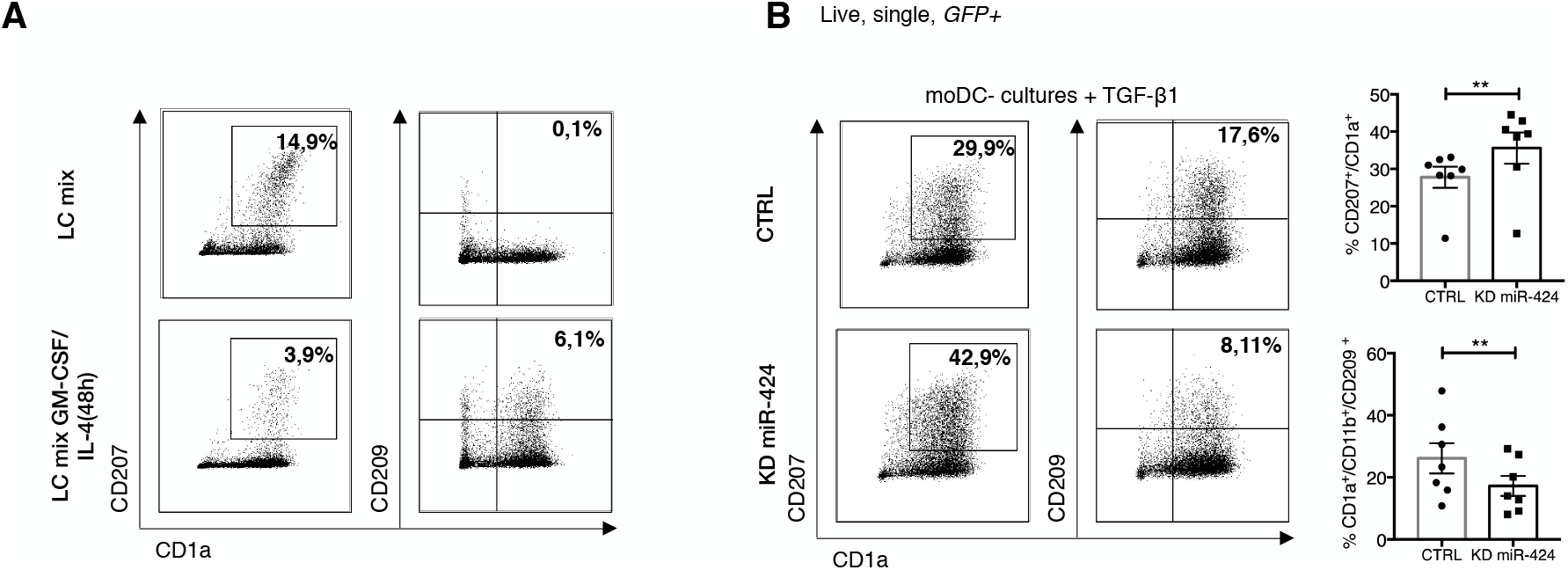
Loss of miR-424 facilitates LC differentiation at the expense of moDC differentiation via TGF-β1 signaling pathway. **A**. Lineage marker profile of human MoPs induced into direct differentiation into LC vs pretreated with GM-CSF/IL-4 for 48hours. **B**. Lineage marker profile of human gated GFP^+^ miR-424 knockdown and CTRL cells, induced into differentiation under lineage specific conditions for 48 hours and then replated into LC-cytokine mix. Data represent the mean ± SEM of 7 donors, 2-tailed Student’s t test, **P<0.01.

**Figure 7.**
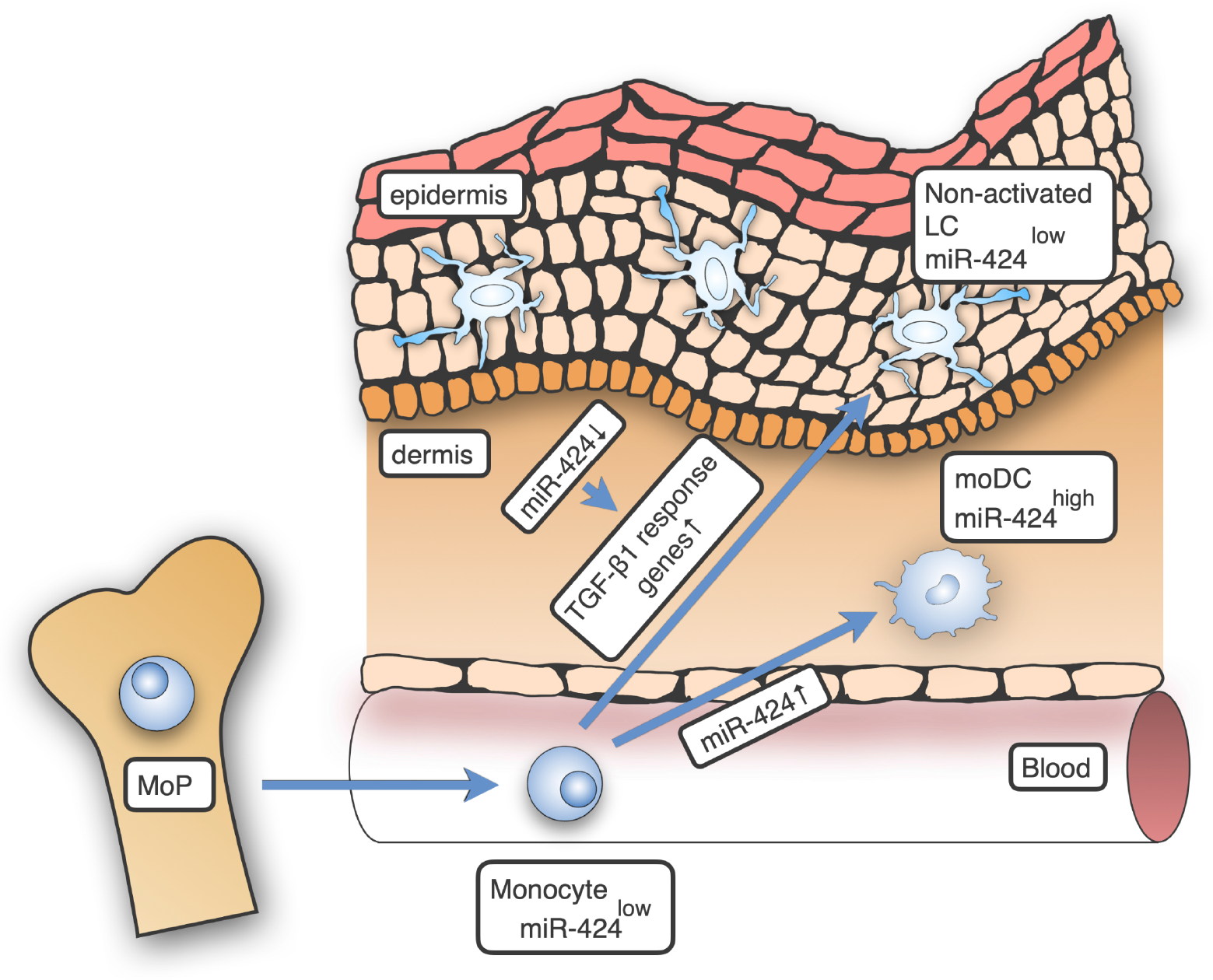
Graphical depiction of the involvement of miR-424(322) in LC vs moDC subset differentiation in the skin by modulating TGF-β signaling.

## Discussion

Murine depletion of candidate DC subsets previously demonstrated that LCs and moDCs exert contrasting functions in skin inflammation. LCs inhibited (Glitzner et al., 2014), whereas moDCs promoted inflammation (Singh et al., 2016). According to a recent concept, these two DC subsets originate from monocyte committed cells (Lutz, Strobl, Schuler, & Romani, 2017). During skin inflammation, both LCs and moDCs were shown to be replenished from peripheral blood monocytes (Collin & Milne, 2016). The intracellular mechanism underlying binary lineage decision of monocytes into LCs vs moDCs is therefore a key switch in the regulation of inflammation.

Using an *in vitro* model of TGF-β1-dpendent LC vs IL-4-dependent moDC differentiation from monocyte intermediates generated in cultures of CD34^+^ human hematopoietic progenitor cells, we here demonstrated that the miR-424/503 cluster is strongly induced downstream of IL-4 concomitant with moDC differentiation, but is repressed during LC differentiation. IL-4 addition is known to inhibit TGF-β1-dependernt LC differentiation (Caux et al., 1999). Consistently, LC differentiation potential is rapidly lost during GM-CSF/IL-4 dependent moDC differentiation. We showed that knockdown of miR-424 during moDC differentiation facilitates TGF-β1-depending LC differentiation, and this effect is paralleled by the induction of TGF-β pathway genes involved in LC differentiation. In line with this, miR-424 knockdown impaired moDC differentiation in human and murine systems, but failed to impair LC differentiation. It even slightly promoted LC numbers *in vivo*. Additionally, LCs from miR-424(322)/503 deficient mice were less activated than LCs from WT mice; both under steady-state and inflammation we observed on average lower numbers of LCs in skin draining lymph nodes. Therefore, low/repressed miR-424 levels during monocyte to LC differentiation facilitates TGF-β1-driven LC differentiation and activation. Thus, together with our observations that miR-424 promotes moDC differentiation, this indicates that miR-424 is a molecular switch regulating anti-inflammatory LC vs pro-inflammatory moDC fates of monocytes. Consistent with this concept, miR-424 is expressed at lower levels by human LCs vs moDCs. This is to our knowledge the first report showing that a miRNA regulates monocyte-derived DC subsets, functionally shown to inversely regulate inflammation.

Bone morphogenetic protein 7 (BMP7) and TGF-β1 are expressed within the epidermis in basal and suprabasal/outer layers, respectively (Yasmin et al., 2013). Moreover, BMP7 is strongly induced in all the keratinocyte layers in the lesional psoriatic epidermis (Borek et al., 2020). We propose a model whereby TGF-β1 ligands repress miR-424 in monocytic LC precursors within the epidermis, which in turn augments TGF-β genes required for TGF-β /BMP7-mediated LC differentiation. Thus, the downregulation of miR-424 may participates in a feed forward signaling loop promoting LC commitment and differentiation within the epidermal microenvironment. Evidence for the existence of such a mechanism is provided by our observations (1) that TGF-β addition to GM-CSF plus TNFα (LC cytokine mix) led to the downregulation of miR-424 concomitant with LC differentiation from monocytic cells; (2) that the miR-424 promoter contains binding sites for RUNX3 potentially repressing miR-424 expression (data not shown); (3) that miR-424 repression during moDC differentiation led to the induction of several genes positively induced in response to TGF-β1-dpendent LC differentiation (i.e. recently described “TGF-β1-LC response genes”). Consequently, the repression of miR-424 may facilitates the expression of TGF-β1-signal-induced anti-inflammatory effector genes such as Axl-1 (Bauer et al., 2012) and miR-146a (Jurkin et al., 2010)

We previously screened for differentially regulated miRNAs during the *in vitro* differentiation of monocytic cells into TGF-β1-dependent LCs vs TGF-β1-independent moDCs (Jurkin et al., 2010)miR-424 stood out in that it was among all 32 miRNAs identified the most strongly inversely regulated one. Additionally, it was co-regulated with miR-503 exhibiting identical target specificity (miR-424/503 cluster) (Jurkin et al., 2010). By applying this semi-functional screen (i.e. TGF-β1 signaling in LC commitment) it was not surprising to identify a miRNA that modulates TGF-β signaling. Consistent with these observations, miR-424 was previously shown to target the TGF-β pathway in epithelial cells (Llobet-Navas et al., 2014). Interestingly, miR-146a was another hit from the same screen, exhibiting opposite regulation relative to miR-424 (i.e. induced in LCs relative to moDCs). miR-146a was expressed at high levels in LCs and miR-424 repressed TLR-dependent NFkB p6/p38-dependent LC activation. However, in contrast to miR-424, miR-146a failed to modulate LC vs moDC differentiation. It will be interesting to further study whether miR-424 repression might promote miR-146a induction during LC commitment and differentiation, towards identifying a gene/miRNA regulatory network securing a non-activated LC state.

It is long known that the addition of IL-4 to LC inducing cytokines represses LC differentiation from monocytic cells, in favor of moDC/dermal DC-like cells (Caux et al., 1999; Göbel et al., 2009; Hoshino et al., 2005; Jurkin et al., 2017). We here showed that IL-4 promotes miR-424 induction during moDC generation within 48h. A key finding of our study was that forced miR-424 repression by lentiviral knockdown facilitated LC differentiation by monocytic cells that have been pre-stimulated with GM-CSF/IL-4. Therefore, miR-424 expression levels seem to represent a molecular switch regulating monocyte differentiation into LC vs moDC. Interestingly, the transcription factor KLF4 is co-regulated with miR-424 during moDC vs LC differentiation (Jurkin et al., 2017). We previously showed that KLF4 has to be repressed to allow LC differentiation and provided evidence that KLF4 transcriptionally represses RUNX3, thereby inhibiting LC differentiation. KLF4 is repressed by Notch signaling (Jurkin et al., 2017)an essential co-signal together with (or downstream promoted by) TGF-β1 for LC differentiation from monocytes (Hoshino et al., 2005; Jurkin et al., 2017) As miR-424 repression in moDC precursors led to the induction of RUNX3, it will be interesting to further study at which levels miR-424 might intersect with KLF4 during the modulation of LC vs moDC differentiation from monocytes. Notably, the miR-424 promoter contains binding sites for TCF4 (data not shown) known to be regulated downstream of Notch/beta catenin signaling (Evans, Chen, Zhang, & Liu, 2010). Thus, it will be interesting to further study the transcription factor network underlying inflammation-associated DC subset differentiation.

Notably, we observed that moDC subsets are reduced in the dermis of miRNA-KO mice under inflammatory conditions in comparison with the WT mice. Interestingly, other immune cell subsets in the skin of miR-KO mice weren’t impaired, suggesting that miR-424 is among DC subsets selectively required for moDC differentiation.

Previous studies identified miR-424 mostly as a tumor suppressive miRNA (Jin, Li, Ouyang, Tan, & Jiang, 2017; Xu et al., 2016), and its function in myeloid cells as well as in human CD34^+^ cells has been poorly understood. Furthermore, miR-424 has previously been functionally linked to TGF-β signaling in epithelial cells: miR-424 is transcriptionally induced by TGF-β1 in the mammary epithelium cells (Llobet-Navas et al., 2014), showing opposite regulation, than here observed in TGF-β1-induced LC differentiation; and it promoted cancer proliferation through targeting TGF-β signaling pathway (Wei et al., 2016). Additionally, opposite regulation by TGF-β1 in epithelial vs LC differentiation has also been shown previously for E-cadherin (Eger et al., 2004; Riedl et al., 2000).

Interestingly, we presented positive regulatory effect of miR-424 in moDC differentiation which seems to be in line with the demonstration of a pro-inflammatory function of miR-424 in the myeloid cell line U937. How miR-424 mediates positive effects on pro-inflammatory moDC differentiation remains to be analyzed further. Among several possibilities, miR-424 might inhibit TGF-β signaling and thereby promote pro-inflammatory characteristics and moDC differentiation; additionally, TGF-β independent effects of genes regulated by GM-CSF might play a positive role in pro-inflammatory cytokine production by moDCs. We identified several candidates (i.e. Tinagl-1, Dntt) to be further analyzed in future studies. miR-424 acted on genes induced by GM-CSF in BM cells, but also acted on genes not regulated by GM-CSF. It will be interesting to further study these groups of genes in moDC differentiation and pro-inflammatory function.

A strength of our study is that we could show both in human and murine models that miR-424 repression facilitates LC differentiation. Thus, our study revealed an evolutionary conserved effect of miR-424 in the regulation of LCs vs moDCs LCs, encouraging future murine clinical translational studies in inflammation and autoimmunity. In comparison, many previous studies pointed to substantial differences between human and murine DC subsets and factors regulating these lineages in human vs mouse. Moreover, miR-424 repression in anti-inflammatory LCs vs induction in pro-inflammatory moDCs might be therapeutically exploited in inflammatory skin diseases. It will be interesting to study whether antagomir-mediated inhibition of miR-424 possess anti-inflammatory pro-TGF-β1-SMAD2/3 activity in autoimmune/inflammatory diseases.

In conclusion, monocyte derived anti-inflammatory vs pro-inflammatory DC fates seem to be inversely regulated by miR-424. MiR-424 is repressed by TGF-β family ligands in monocytic LC precursors within epithelia, and miR-424 repression may facilitate LC differentiation by augmenting the expression of members of the TGF-β family pathway. Moreover, miR-424 repression seems to be involved in securing a high activation threshold in LCs.

## Materials and methods

### Cell isolation and *in vitro* differentiation

Human cord-blood CD34^+^ cells were isolated by magnetic sorting using EasySep human CD34 positive selection kit (Stem Cell Technologies) according to manufacturer’s protocol. Cord blood was collected during healthy, full-term deliveries. CD34^+^ cord-blood cells were cultured for 2-3 days under progenitor expansion conditions (serum free X-Vivo media, 50 ng/ml SCF, 50 ng/ml FLT3L and 50 ng/ml TPO). For direct LC differentiation CD34^+^ cells were cultured for 7 days in a 24 well tissue culture plate (5×10^4^ cells per well) in serum free CellGro DC medium (CellGenix, Freiburg, Germany) supplemented with Glutamax, penicillin/streptomycin, 2.5 ng/ml TNFα, 100 ng/ml, GM-CSF, 50 ng/ml FLT3L, 20 ng/ml SCF and 1 ng/ml TGF-β1. For direct moDC differentiation CD14^+^ monocytes were isolated from PBMC by positive selection using Miltenyi Biotec kit. Then CD14^+^ monocytes were differentiated in RPMI medium (Sigma, St. Luis, Mo, USA) supplemented with 10% FBS (Sigma-Aldrich, St. Louis, Mo, USA), 100 ng/ml GM-CSF, and 35 ng/ml IL-4. For indirect moDCs and LCs differentiation previously described two-step culture model was applied with slight modifications (Caux et al., 1997; Jurkin et al., 2010). Briefly, CD34^+^ cells were plated (5 × 10^4^ to 1× 10^5^ / ml per well) in 24-well tissue culture plate in CellGro DC medium supplemented with 10% FCS, 100 ng/mL GM-CSF, 20 ng/mL SCF, 50 ng/mL FMS-FLT3L, and 2.5 ng/mL TNF-α for 3-5 days and subsequent differentiation in RPMI (Sigma, St Louis, Mo; +10% FCS) either with 100 ng/mL GM-CSF, 35 ng/mL IL-4 for moDCs or100 ng/mL GM-CSF, 2.5 ng/mL TNF-α, and 1 ng/mL TGF-β1 for LCs for 5-6 days. To generate moMacropages, CD14^+^ monocytes isolated from buffy coat were differentiated in RPMI medium supplemented with 100 ng/mL GM-CSF and 2 ng/mL M-CSF in the presence of 10% FCS for 6-7 days. For moDCs generation from CD14^+^ monocytes, cells were cultured for 6 days in RPMI medium +10% FCS either with addition of 100 ng/mL GM-CSF and 35 ng/mL IL-4 or 100 ng/mL GM-CSF alone. Granulocytes were generated from CD34^+^ cord-blood cells by plating 300.000 cells in FCS free RPMI medium supplemented with 100 μg/mL G-CSF and 20 μg/mL SCF. On day 5 and 8 cells were replated in fresh medium and harvested on day 10 of differentiation. All cell cultures were supplemented with GlutaMAX (2.5 mmol/L; Invitrogen, Grand Island, NY) and penicillin/streptomycin (125 U/ml each; PAA, Pasching, Austria). Neutrophils were isolated from peripheral blood by density-gradient centrifugation using Lymphoprep (Axis-Shield PoC AS, Oslo, Norway) according to the manufacturer’s protocol.

### Cytokines and reagents

Cytokines tumor necrosis factor α (TNFα), thrombopoietin (TPO), stem cell factor (SCF), Fms-like tyrosine kinase 3 ligand (FLT3-L), granulocyte-macrophage colony stimulating factor (GM-CSF), interleukin-4 (IL-4) were purchased from PeproTech, UK. Human recombinant transforming growth factor-β1 (TGF-β1) was purchased from R&D, USA.

### Lentiviral transfection, transduction of CD 34^+^ cord-blood cells

To silence miR-424 the GFP tagged lentiviral vectors that express antisense hsa-miR-424 (MZIP424-PA-1) and corresponding scramble control lentiviral vectors (MZIP000-PA-1) were purchased from SBI (System Biosciences). Transfection of 293T packaging cell line was performed using calcium-phosphate protocol and infection of target cells was done as previously described (Platzer et al., 2004). In general, lentiviral supernatant was added to CD34^+^ cells in the RetroNectin (Takara) precoated 24-well suspension plate according to manufacturer protocol. Infections were repeated 2-3 times and on day 5 cells were replated into lineage-specific cytokine mix and further differentiated for 5-7 days as previously described. Then cells were harvested, stained and acquired to GFP^+^ gate.

### Experimental animals and treatment

Animal maintenance and experiments were performed under the Institutional Animal Care and Use Committee (IACUC) guidelines. Protocol number is IACUC-2018-0008. Experiments were performed with miR-424(322) ^−/−^ male mice with homogeneous FvB/NJ background. For genotyping hair samples were incubated in 250 μL of Direct PCR reagent and 20 mg/mL Proteinase K (New England Biolabs, P8102) for 1 h at 55°C. PCR was performed using FastStart polymerase (Roche, 047384200010 under the following conditions: predenaturation for 5 min at 94°C, 35× (denaturing for 30 sec at 94°C, annealing for 30 sec at 55°C, and elongation for 30 sec at 72°C), and final elongation for 7 min at 72°C.

Mice 8-14 weeks old were topical applied of 62 mg of 5% Aldara IIMQ cream as previously was determined as the most optimal dose to induce ear inflammation(van der Fits et al., 2009). Mice were treated with IMQ daily at the same time for 1 week and ear thickness was measured daily using electronic measurement device C1X018 (Kroeplin, GmbH).

### Purification and generation of mouse BM-derived dendritic cells

BM-derived LCs and moDCs were generated *in vitro* from mouse BM precursors. In brief, femur and tibiae were removed from 8-12-week-old mice. BM cells were flushed out with ice-cold PBS, washed and incubated in 1 ml Red Blood Cell Lysing Buffer (Hybri-Max, Sigma-Aldrich) for 1 minute. After washing cells were plated in 24-well tissue culture plate (0.5 × 10^6^ /ml per well) in DMEM supplemented with 1% penicillin/streptomycin (Lonza #CC-4136), 10% FCS and 1% MEM non-essential amino acids solution (ThermoFisher) under lineage specific cytokine condition for LCs (20 ng/ml GM-CSF, 1 ng/ml TGF-β1) and moDCs (20 ng/ml GM-CSF). On day 3, the medium was changed and cells were harvested on day 7.

### Flow cytometry

All flow cytometry experiments were performed on LSRII and LSR Fortessa flow cytometers (BD Biosciences, USA). The data was analysed using the DIVA (BD Biosciences) and FlowJo software (Tree Star, Inc.USA). For FACS sorting the FACS Aria instrument (BD Biosciences) was used. Human cells were harvested, resuspended in 40 μl of staining buffer and stained on ice for at least 1 hour. Fc receptors were blocked by incubating for 15 minutes on ice before staining with antibodies. Mouse ears were harvested and separated for ventral and dorsal sheets. The sheets were incubated for 1 hour at 37□°C in 0.8% Trypsin solution. Then epidermal and dermal sheets were digested separately as previously described(Singh et al., 2016). Single cell suspension was incubated for 1:1000 diluted purified anti-mouse CD16/32 (Biolegend) for at least 20 min on ice before adding antibodies. Dead cells were excluded by using Zombie NIR viability dye (Biolegend). All the flow cytometry antibodies were summarized in the Table S2.

### RNA-isolation and real time PCR

The extraction of RNA was performed using miRNAeasy Mini Kit (Qiagen) with DNase I treatment according to the manufacturer protocol. RT-PCR was performed using TaqManTM MicroRNA Assay. Probes for TaqManTM MicroRNA Assay (hsa-miR-424-3p, hsa-miR-424-5p, U6 snRNA control) were purchased from ThermoFisher. All the steps were performed according to manufacturer’s instructions.

### Immunohistochemistry on mouse epidermal sheets

Epidermal sheets of mouse ears were fixed in 4% PFA for 30 minutes and incubated for 2 hours in blocking solution (1% BSA, 0,05% horse serum, 20%Triton in PBS) at RT. Staining was performed using 1:1500 dilution Hoechst 33342 (ThermoFisher), 1:500 diluted MHCII/I-A/I-E PE (M5/114.15.2, Biolegend) and 1:200 diluted CD207 AF488 (929F3.01, Acris Antibodies, USA) FACS antibodies. Epidermal sheets were washed twice with PBS and analysed on the Zeiss Axio Obzerver Z1 microscope. Images were processed using ImageJ software.

### Epidermal thickness measurement

Mouse ears were fixed in formalin for 1-2 days. After paraffin embedding the sections were stained H&E (Sigma, USA) and processed at Leica DM4000 B microscope (Leica Cambridge Ltd) equipped with Leica DFC320 Video camera (Leica Cambridge Ltd). Epidermal thickness (10X magnification) was measured in 10 random fields on 3 pictures per sample. Analysis was performed using ImageJ software.

### mRNA sequencing and data analysis

Cells for total RNA sequencing were collected after extraction from mouse BM and 24h after addition of GM-SCF and frozen in lysis buffer until RNA extraction. Total RNA was isolated using either RNeasy Micro Kit (Qiagen) or miRVana isolation kit (Ambion). RNA quality was checked at Bioanalyser before library preparation. Stranded RNA libraries were prepared with 100ng total RNA using the KAPA RNA HyperPrep Kit with RiboErase (HMR) (Roche). RNA sequencing was performed on Illumina NovaSeq 6000 system (Illumina). Gene expression was quantified in counts using Salmon. Each data point has 3 biological replicates. In both day 0 and day 2, differential expression between miR-KO and wild type was estimated using DESeq2. Differential expressed genes were selected based on the adjusted P-value with cut off equal 0.05. The RNA-seq data will be deposited in the NCBI Gene Expression Omnibus (GEO).

### Statistical analysis

Statistical analysis was carried out using GraphPad Prism software. Student’s t-test was used for comparing differences between two groups. Multiple groups were subjected to analysis of variance (ANOVA) analysis. P values <0.05 were considered statistically significant.

## Abbreviations used

BM: bone marrow
BMDC: bone marrow-derived dendritic cells
BMP7: bone morphogenetic protein 7
DC: dendritic cell
GM-SCF: granulocyte-macrophage colony-stimulating factor
IL-4: interleukin 4
IMQ: imiquimod
KLF4: Kruppel-like factor 4
LC: Langerhans cell
miR-424: microRNA-424
moDC: monocyte-derived dendritic cell
moPs: monocyte committed progenitor cells
RUNX3: runt-related transcription factor 3
TGF-β1: Transforming growth factor beta 1

## Online supplemental material

Fig. S1 contains the analysis of miR-424 role in human dendritic cell differentiation *in vitro*.

Fig. S2 shows the Immune cell phenotyping in WT vs miR-KO mice upon IMQ-induced inflammation.

**Table S1.** shows the list of TGF-β1 response genes upregulated in miR-424 knockout BMDCs.

|  | Gene | 6h.logFC | 6h.P.Value | 24h.logFC | 24h.P.Value |
| --- | --- | --- | --- | --- | --- |
| 1 | OLR1 | 4,66 | 8,16E-07 | 5,83 | 5,16E-07 |
| 2 | SKIL | 3,11 | 2,27E-04 | 3,26 | 1,19E-04 |
| 3 | RGS16 | 2,99 | 3,06E-04 | 3,56 | 4,21E-05 |
| 4 | VASH1 | 2,49 | 1,57E-05 | 2,77 | 2,22E-05 |
| 5 | DOCK4 | 2,36 | 1,51E-04 | 3,02 | 5,14E-04 |
| 6 | RGS1 | 2,35 | 5,54E-05 | 2,99 | 1,33E-05 |
| 7 | PLXDC2 | 2,32 | 4,23E-04 | 2,62 | 5,11E-05 |
| 8 | CDH1 | 2,20 | 7,81E-04 | 4,45 | 8,31E-06 |
| 9 | TNFSF14 | 2,19 | 3,59E-04 | 1,97 | 1,50E-04 |
| 10 | FMNL3 | 1,81 | 5,38E-04 | 1,46 | 5,42E-04 |
| 11 | MS4A7 | 1,70 | 5,39E-04 | 2,05 | 1,04E-04 |
| 12 | TRIM36 | 1,67 | 9,38E-04 | 1,82 | 1,77E-04 |
| 13 | MMP2 | 1,57 | 4,72E-04 | 1,60 | 6,92E-04 |
| 14 | CD9 | 1,51 | 8,91E-04 | 2,65 | 8,37E-05 |
| 15 | HAMP | 1,51 | 1,25E-03 | 2,58 | 6,23E-04 |
| 16 | PDGFA | 1,45 | 2,01E-03 | 2,42 | 4,83E-05 |
| 17 | FAM43A | 1,41 | 1,63E-03 | 1,57 | 9,84E-04 |
| 18 | CDH26 | 1,34 | 8,31E-04 | 1,74 | 2,60E-04 |
| 19 | ID2 | 1,33 | 6,90E-04 | 1,62 | 3,47E-04 |
| 20 | RUNX3 | 1,29 | 1,99E-04 | 1,28 | 3,98E-04 |
| 21 | TGFBI | 1,24 | 1,70E-02 | 2,40 | 7,24E-04 |
| 22 | PROCR | 1,20 | 5,63E-03 | 1,87 | 1,25E-04 |
| 23 | P2RY13 | 1,06 | 6,57E-03 | 1,49 | 6,05E-04 |
| 24 | SNX7 | 1,03 | 1,90E-03 | 1,99 | 3,32E-04 |
| 25 | RASGRP3 | 1,00 | 1,40E-02 | 1,56 | 7,88E-04 |
| 26 | LAPTM5 | 0,75 | 4,84E-04 | 1,03 | 8,98E-05 |
| 27 | STX6 | 0,65 | 2,48E-02 | 1,31 | 9,56E-04 |
| 28 | B3GNT7 | 0,60 | 2,59E-01 | 1,85 | 7,85E-04 |
| 29 | ST14 | 0,60 | 7,71E-02 | 1,74 | 2,54E-04 |
| 30 | IL18 | 0,58 | 2,80E-01 | 1,84 | 8,27E-04 |
| 31 | LCP2 | 0,57 | 3,11E-02 | 1,41 | 4,05E-04 |
| 32 | SH3BP4 | 0,40 | 1,86E-01 | 1,34 | 9,14E-04 |
| 33 | LAYN | 0,38 | 1,61E-01 | 1,56 | 7,34E-04 |

**Table S2.**
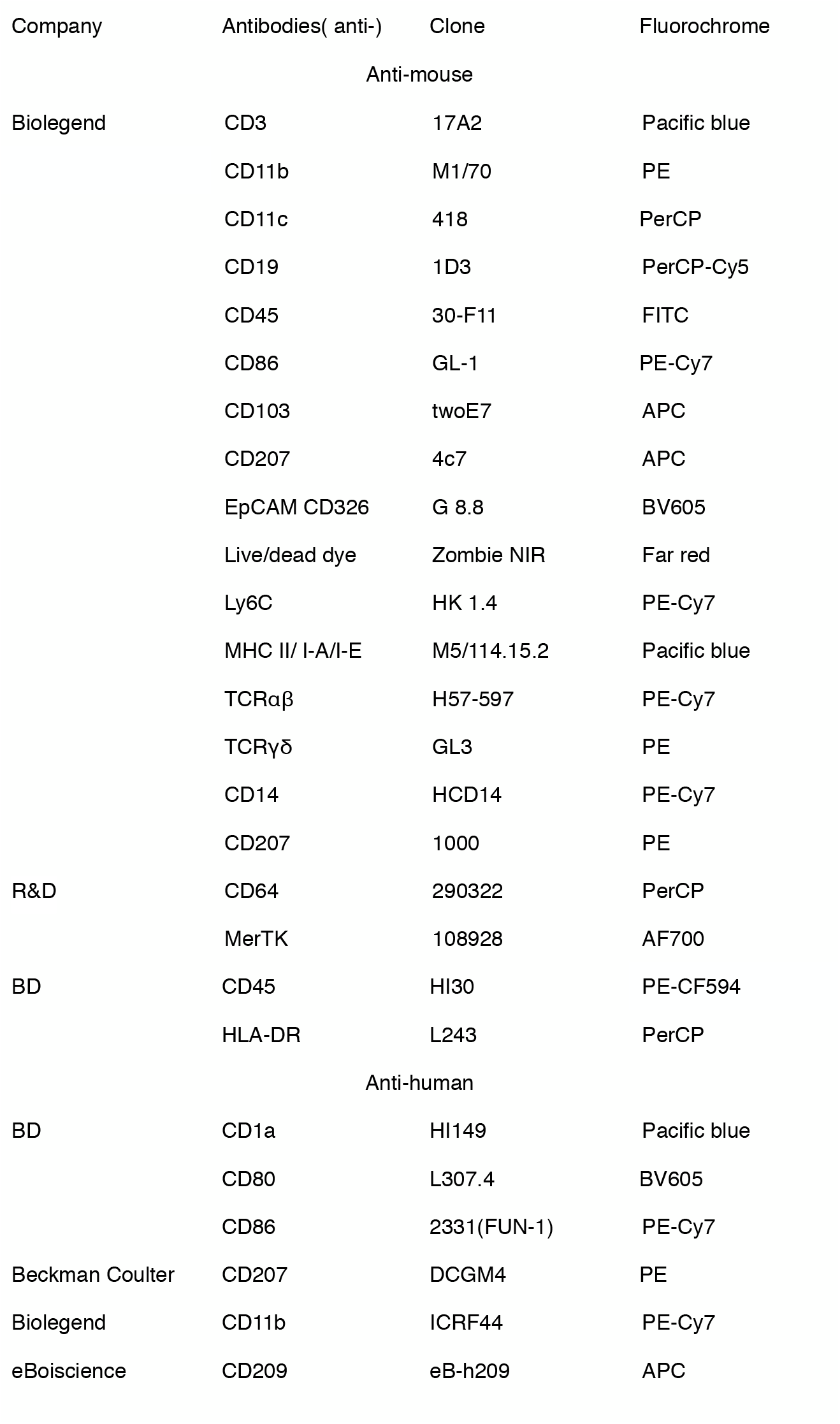
lists the flow cytometry antibodies.

## Acknowledgements

This work was supported by the Austrian Science Fund FWF (W1241), the PhD program Molecular Fundamentals of Inflammation (DK-MOLIN) of the Medical University of Graz and the Austrian Marshall Plan Foundation. We thank the flow cytometry core facility of the Icahn School of Medicine at Mount Sinai (ISMMS), Erin Nekritz (ISMMS) for mouse breeding and Elke Schwarzenberger (MUG) for cell preparation and technical assistance.

## Declaration of interests

The authors declare that they have no relevant conflicts of interest.

## Author contributions

V.Z and H.S. contributed to the design and conception of the study, analyzed and interpreted the data, co-wrote the manuscript: C.Passegger. T.S., C.Pollack, A.Z. contributed to data acquisition, analysis and experimental design; C.T. performed histology staining; K.K.,B.J.,Q.P.,B.S.,J.E,J.Y performed RNA-sequencing and carried out bioinformatical analysis; J.M.S. provided miR-KO mice, contributed to experimental design and data interpretation, revised the manuscript. All authors have read and approved the final manuscript.

## Supplementart=y figure legends

**Figure S1.**
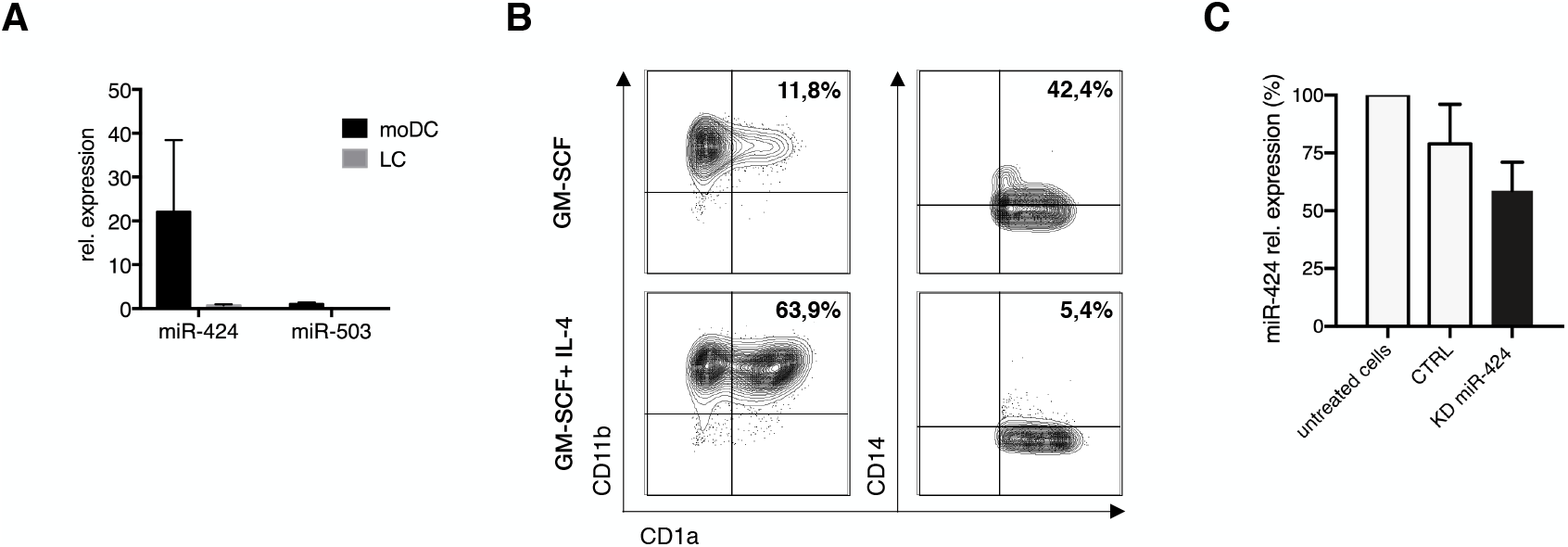
Analysis of miR-424 role in human dendritic cell differentiation *in vitro*. **A**. Relative expression of miR-424- and miR-503 measured by quantitative Taqman PCR (n=3, ± SEM). **B**. Phenotype of moDCs differentiated from CD14^+^ monocytes with GM-CSF+IL-4 or GM-CSF alone. **C**. Knockdown of miR-424 in transduced monocytic precursors and untreated cells harvested on the 5^th^ day of differentiation from CD34^+^ cells. Transduced cells were >85% GFP^+^ according to FACS (n=8, ± SEM).

**Figure S2.**
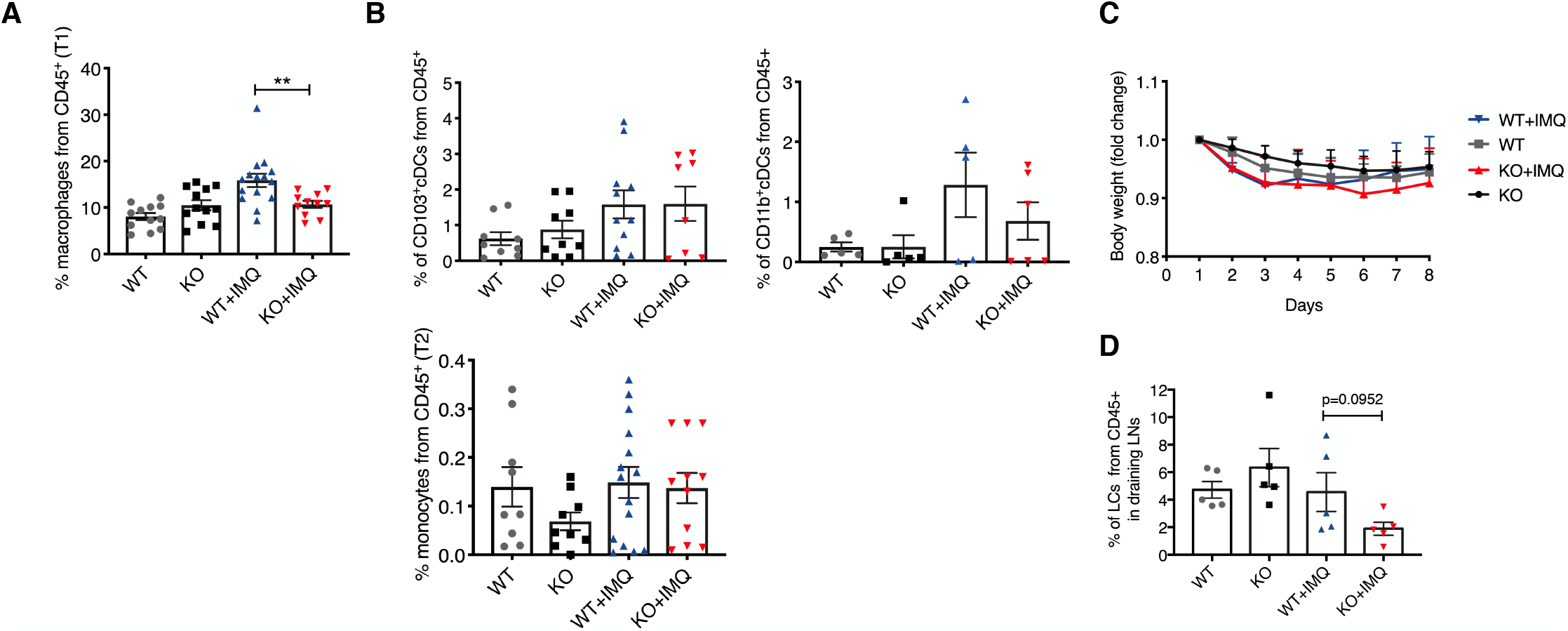
Immune cell phenotyping in WT vs miR-KO mice upon IMQ-induced inflammation. **A**. Graph represents percentage of phenotypically defined macrophages (T1) and monocytes (T2) m in mouse dermis quantified by flow cytometry on day 7 of experiment (n=10-12, ± SEM, unpaired Student’s t test, ** P<0.01). **B**. Graph represents percentage of phenotypically defined CD103^+^ cDCs, CD11b^+^ cDCs and monocytes (T2) in mouse dermis quantified by flow cytometry on day 7 of experiment (n=6-10, ± SEM, unpaired Student’s t test). **C**. Body weight in WT and miR-424 KO mice on the indicated days (n=6-8 mice per group, ± SEM). **D**. Graph shows frequency of LCs in draining lymph nodes on day 7 of IMQ experiment.

## Notes

### Competing Interest Statement

The authors have declared no competing interest.

